# Integrative reverse vaccinology approach for identification of a vaccine antigen candidate with immunoprotective and anti-fibrotic potential against *Schistosoma japonicum*-associated liver fibrosis

**DOI:** 10.1101/2025.09.02.673671

**Authors:** Qian Xu Yang, Jue Wang, Ying Zi Ming, Yu Zhang, Li Ping Wong, Hai Yen Lee

## Abstract

*Schistosoma japonicum-*associated liver fibrosis (SSLF) is a major cause of morbidity in Schistosomiasis, yet no licensed vaccines or specific therapies exist to prevent or treat this complication. Current vaccine development has focused mainly on infection prevention, with limited attention to pathology-driven outcomes such as fibrosis. To address this gap, we developed an in-silico pipeline integrating host single-cell transcriptomic analysis, parasite antigen screening, and structure-based modeling. Single-cell RNA-seq of liver tissues from SSLF and controls identified three fibrosis-related hub proteins: FYN, BCL2, and AKT3. In parallel, parasite antigens curated from public databases were evaluated for immunogenicity and safety using reverse vaccinology principles. Among these, DRE2_SCHJA, an Anamorsin homolog involved in Fe–S cluster assembly, was prioritized as the top candidate. Docking analysis predicted the strongest interaction between DRE2_SCHJA and FYN (ΔG = −14.9 kcal/mol), linking the antigen to a central regulator of fibrosis. Unlike Saracatinib, which inhibits FYN at the ATP–binding pocket, DRE2_SCHJA bound to the SH2 domain through both canonical and non-canonical contacts, indicating a different regulatory mechanism. Taken together, these findings suggest DRE2_SCHJA as a potential vaccine candidate with both protective and anti-fibrotic potential against SSLF. While experimental validation is required, this study supports the early identification of SSLF vaccine candidates in a rapid and cost-effective manner and provides a framework applicable to other neglected tropical diseases.

**Athor summary:** *Schistosoma japonicum (S. japonicum)* is a parasitic disease that remains a major health problem in parts of Asia, especially in China. One of its most harmful consequences is liver fibrosis, which causes long-term illness and reduces quality of life. Currently, there are no drugs or vaccines that specifically prevent or treat this condition. In this study, we used a fast and low-cost computer-based method to search for vaccine targets against liver fibrosis caused by *S. japonicum*. We identified a parasite protein that could serve as a vaccine target, with the potential to prevent fibrosis from developing and to slow its progression. If future studies confirm its role, such a vaccine could greatly improve the long-term health of people living with schistosomiasis and inspire new strategies for vaccine development in other neglected tropical diseases.

## Introduction

Schistosomiasis remains one of the most devastating negelected tropical diseases, second only to malaria in terms of disease burden, affecting over 240 million people and placing more than 700 million individuals at risk in endemic regions[1]. Among human schistosomes, *Schistosoma japonicum (S. japonicum)* is mainly endemic to East Asia, particularly China. Compared with other species, *S. japonicum* infection is characterized by high egg output and strong invasive capacity [2]. The most severe consequence of chronic *S. japonicum* infection is *Schistosoma japonicum*-associated liver fibrosis(SSLF). It develops when parasite eggs become trapped in the hepatic portal system, where they provoke persistent granulomatous inflammation and drive progressive fibrogenesis[3, 4]. Without timely intervention, liver fibrosis can progress to cirrhosis, liver failure, or hepatocellular carcinoma[5, 6], which is a major contributor to long-term morbidity and mortality in endemic populations.

Currently, praziquantel (PZQ) is the only widely used drug for Schistosomiasis chemotherapy, administered mainly through large-scale mass drug administration (MDA) programs. However, the long-term effectiveness of this strategy has been limited by persistent challenges, including insufficient drug delivery, poor adherence, high reinfection rates, and inadequate health infrastructure in endemic regions. Although PZQ is highly effective against adult worms, it has no activity against juvenile schistosomes and does not prevent reinfection, leading to continuous egg deposition and further accelerating SSLF[7]. Moreover, it has no effect on parasite eggs already lodged in host tissues, which remain the principal drivers of chronic inflammation and fibrogenesis[8]. In addition, PZQ lacks intrinsic antifibrotic activity, meaning that established liver fibrosis continues to advance even after complete clearance of adult worms[9]. In recognition of these limitations, China’s National Plan for Schistosomiasis Elimination (2023–2030) emphasizes the prevention and treatment of SSLF and calls for the development of novel strategies, including vaccines, to complement existing control measures and mitigate long-term morbidity.

Despite these challenges, Schistosomiasis vaccine development remains slow, and it is considered among the top ten most urgently needed vaccines worldwide[10]. To date, only four vaccine candidates—Sh28GST (phase III), Sm14 (phase II), Sm-TSP-2 (phase II), and Sm-p80 (phase I)— have progressed to clinical trials. Progress has been hampered, with Bilhvax (Sh28GST) failing to show efficacy in reducing Schistosomiasis recurrence in children, despite inducing immune responses dominated by non-protective antibody subclasses[11, 12]. Importantly, most of these efforts have focused on *Schistosoma mansoni*, leaving vaccine development for *S. japonicum* comparatively underexplored. The latest review about the Schistosomiasis vaccine has emphasized that the vaccine research should not be limited to infection prevention but should also address chronic complications[13].

Recent advances in high-throughput technologies and computational immunology have provided new opportunities to overcome persistent barriers in Schistosomiasis vaccine development[14]. Reverse vaccinology (RV)-based antigen prediction enables the rapid, in silico evaluation of candidate immunogenicity and safety, thereby reducing reliance on early stage animal experiments and lowering development costs[15]. Inspired by this approach, we applied an intergrative in silico strategy to screen potential antigen associated with SSLF. Recognizing that parasite–host interactions shape pathological processes[16, 17], yet their integration into vaccine research remains largely unexplored. To extend the preventive function of vaccine targets, we investigated whether candidate antigens could also modulate SSLF pathology. Using single-cell transcriptomic analysis of host liver tissue, we identified key fibrosis-related proteins as potential host targets. Structural docking was then performed to assess antigen–protein interactions, and these binding patterns were compared with those of a known antifibrotic inhibitor acting on the same hub protein. These approaches allowed us to suggest candidate antigens that may elicit protective immunity while potentially modulating fibrosis progression. This strategy offers a new paradigm for Schistosomiasis vaccine development and may also inform approaches for other NTDs.

## Materials and method

### Identification of hub gene for SSLF

The workflow for identifying SSLF-associated hub genes is outlined in Fig 1, integrating single-cell transcriptomics, co-expression network, differential expression, and PPI analyses[18, 19].

**Fig 1.**
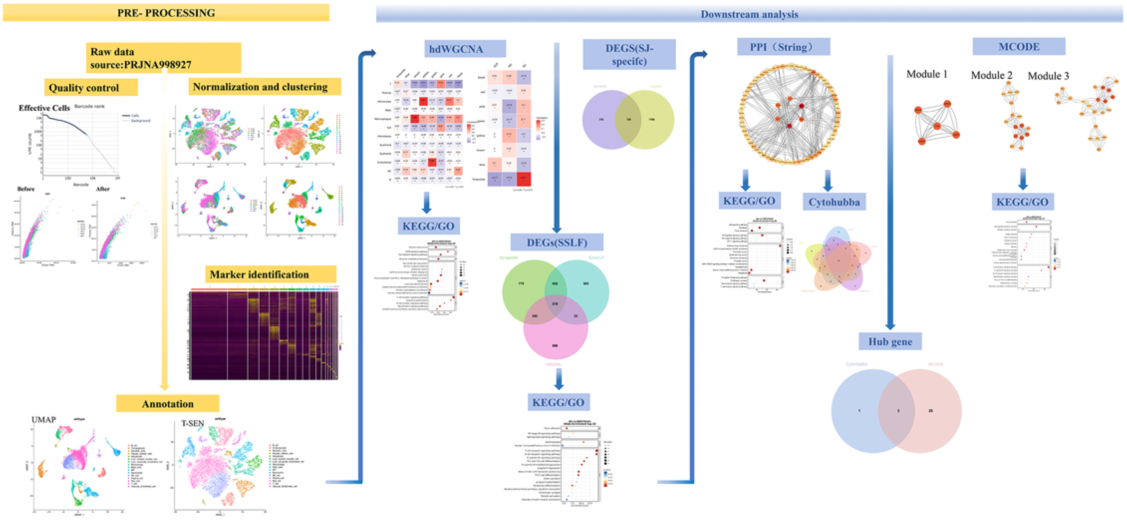
Workflow of single-cell RNA analysis and hub gene identification in *Schistosoma japonicum*-associated liver fibrosis

### Single-cell RNA sequencing analysis

Single-cell RNA sequencing (scRNA-seq) data were obtained from the PRJNA998927 dataset, originally generated by Yu Zhang *et al.* (2021), comprising liver tissues from *S. japonicum*–HBV co-infection (SJ, n = 1), HBV-induced fibrosis (CLF, n = 3), and healthy controls (HC, n = 2). Raw reads were processed with CeleScope (v2.0.7) for barcode filtering and UMI deduplication and aligned using STAR (v2.7.3a) [20]. Cells were filtered in Seurat (v4.3.0) based on gene count (200–10,000) and mitochondrial content (< 25%) [19], with doublets removed by DoubletFinder (v2.0.3). Normalization and identification of highly variable genes were followed by PCA, clustering, and dimensionality reduction (UMAP/t-SNE). Cell types were annotated using SingleR (v2.0.0) and refined by manual inspection of cluster-specific marker genes, with highly similar clusters merged for downstream analysis.

### High-dimensional weighted gene co-expression network analysis (HD-WGCNA)

Gene co-expression modules were identified using HD-WGCNA (v0.3.01) on a Harmony-integrated Seurat object[21]. Genes expressed in ≥ 5% of cells were retained, and metacells were constructed per cell type and condition using k-nearest neighbors (k = 25, max_shared = 10) on Harmony-reduced dimensions. A signed hybrid network was built with a soft-thresholding power of 3 (scale-free R² ≈ 0.95, mean connectivity ≈ 30). Modules were detected by hierarchical clustering of the TOM and refined with dynamic tree cutting. Module eigengenes (MEs) and memberships (kME) were calculated, and module activity was assessed by UCell. Module-trait associations with condition (HC, CLF, SJ) and cell types were determined using Pearson correlation (p < 0.05). Significant modules were subjected to GO and KEGG enrichment using clusterProfiler.

### Differentially expressed genes (DEGs)

Differential expression was analyzed with Seurat’s FindMarkers (adjusted p < 0.05, |log₂FC| ≥ 1) [22] across SJ vs. HC, CLF vs. HC, and SJ vs. CLF. SSLF-specific genes were defined as DEGs in SJ vs. HC but not CLF vs. HC, then intersected with the DEGs of SJ vs. CLF and the WGCNA turquoise module. Then, these genes were subjected to GO and KEGG enrichment.

### Protein-protein interaction network (PPI) construction and module analysis

A PPI network was constructed using STRING (score > 0.4) and visualized in Cytoscape (v3.10.3). Hub genes were identified by five CytoHubba algorithms (MCC, MNC, Degree, Betweenness, Closeness) and MCODE clustering (default parameters). Genes intersecting the top ten CytoHubba rankings and top three significant MCODE modules were defined as SSLF hub genes[23].

### Identification of potential antigens against SSLF

The antigen screening workflow (Fig 2) included filtering reviewed *S. japonicum* proteins by GO annotations, structural features, and host similarity, followed by B-cell, CTL, and HTL epitope prediction. Peptides meeting both immunogenicity and safety criteria were retained as SSLF vaccine candidates.

**Fig 2.**
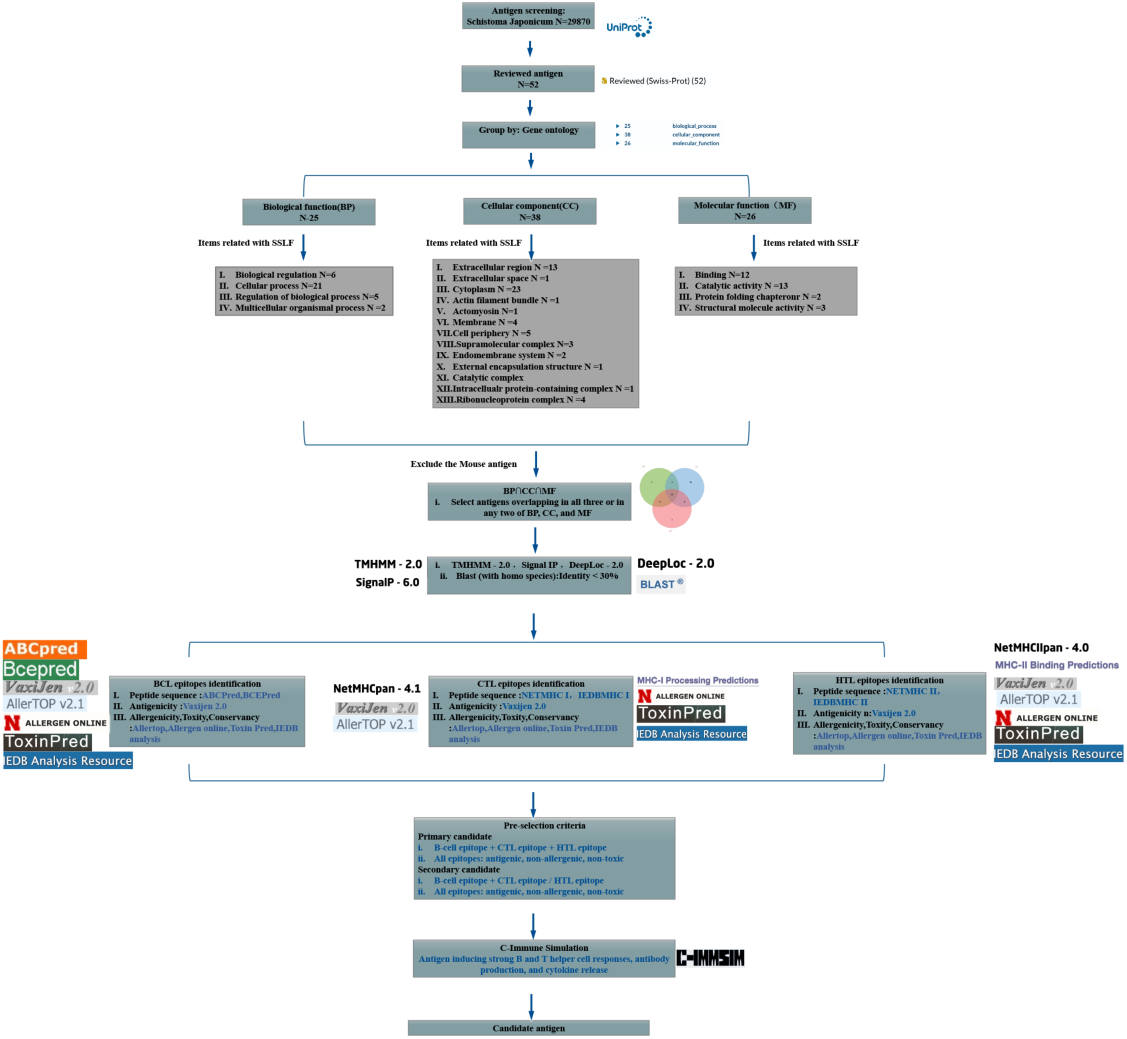
Antigen screening workflow to identify vaccine candidates against *Schistosoma japonicum*-associated liver fibrosis

### Antigen selection based on GO annotation and homology exclusion

Reviewed *S. japonicum* proteins were retrieved from UniProt and filtered by Gene Ontology (GO) terms related to liver fibrosis[24, 25]. Antigens annotated in at least two GO domains (Biological Process, Cellular Component, Molecular Function) were retained. Structural features and localization were predicted using TMHMM, SignaIP, and DeepLoc, and the results were used for reference rather than as exclusion criteria. Proteins showing ≧ 30% identity to human sequences (BLAST) were excluded[26].

### BCL epitope prediction

Linear B-cell epitopes were predicted using ABCPred and BCEPRED under default settings. ABCPred candidates with scores ≥ 0.85 were retained, while BCEPRED predictions were selected based on highlighted sequences. Peptides identified by both tools were evaluated for antigenicity using VaxiJen v2.0[27].

### CTL epitope prediction

Cytotoxic T lymphocyte (CTL) epitopes were predicted using NetMHCpan-4.1 and the IEDB MHC-I binding tool. In NetMHCpan-4.1, peptides classified as strong binders (SB) were selected; weak binders (WB) were included only when SBs were absent. Peptides with a binding rank ≤1% in IEDB were also retained. Only peptides predicted by both tools were assessed for antigenicity via VaxiJen v2.0[27].

### HTL epitope prediction

Helper T lymphocyte (HTL) epitopes were identified using NetMHCII and IEDB MHC-II tools. Selection followed the same criteria as CTL prediction: SB or WB from NetMHCII and ≤ 1% binding rank from IEDB. Epitopes predicted by both were evaluated for antigenicity using VaxiJen v2.0[27].

### Allergy, toxicity and conservancy prediction

Allergenicity was assessed with AllerTOP 2.0 and AllergenOnline, toxicity with ToxinPred3.0, and conservancy with the IEDB Epitope Conservancy Tool[26].

### Antigen selection criteria

Antigens were defined as primary candidates if they contained at least one B-cell epitope and both CTL and HTL epitopes, all showing antigenicity, non-allergenicity by both allergenicity tools, non-toxicity, and 100% conservancy. Antigens meeting these criteria but containing only one type of T-cell epitope were considered secondary candidates[28]. This step ensured that only strongly immunogenic antigens advanced to further validation[29, 30].

### Immune simulation of antigens

In silico immune simulations of candidate antigens were performed using C-ImmSim (default parameters)[31]. The simulation volume was 10, random seed 12345, and 100 time steps, with a single injection at step 1. HLA alleles prevalent in the Chinese population were selected[32]. Antigens inducing strong B cell activation, helper T cell responses, sustained antibody production, and cytokine release were retained as final vaccine candidates.

### Docking and computational analyses

#### First protein–protein docking

Protein-protein docking was performed to assess interactions between *S. japonicum* antigens and host fibrosis-related hub proteins. Structures were obtained from UniProt, prioritizing high-resolution data or AlphaFold models. Non-protein molecules were removed in PyMOL (v2.5.3). Docking was conducted using ClusPro (default parameters)[33], with complexes ranked by ClusPro scores and binding affinities predicted by PRODIGY. Top complexes were examined in PyMOL for interfacial interactions (hydrogen bonds, salt bridges, π–π stacking), and final evaluation was based on predicted binding free energy (ΔG) supported by docking scores and interface features.

#### Molecular docking

Molecular docking was performed in Maestro (v13.9, Schrödinger LLC) to validate the structural reliability of the selected hub protein[34]. The protein was prepared with the Protein Preparation Wizard (hydrogen addition, bond order assignment, H-bond optimization, and restrained minimization using the OPLS4 force field). A clinically approved inhibitor was prepared using LigPrep to generate low-energy conformations and appropriate protonation states at physiological pH (7.0 ± 0.5). Docking was conducted with Glide in extra precision (XP) mode. Poses were ranked by docking score, and the top-ranked pose was used to confirm structural suitability for subsequent analyses.

#### Secondary protein–protein docking

Because the hub protein structure used in the initial docking differed from the conformation targeted by its known inhibitor, protein-protein docking was repeated using the antigen and the inhibitor-validated hub protein structure from the previous molecular docking step. The same ClusPro workflow was applied, with the aim of comparing the antigen-protein binding interface against the reference site defined by the inhibitor–protein complex.

#### Interface visualization

Docking results were visualized in PyMOL (v2.5.3) by aligning the hub protein structure and showing the binding sites of both the inhibitor and the antigen to compare their spatial distribution[35].

#### Motif prediction

To explore antigen–host protein interactions, short linear motifs (SLiMs) in the antigen sequence were predicted using the Eukaryotic Linear Motif (ELM) resource (http://elm.eu.org) to identify functional sites relevant to protein binding and regulation[36].

## Results

### Single-cell RNA sequencing analysis

Single-cell RNA sequencing was performed on six liver samples, generating high-quality data with valid read rates exceeding 96% and Q30 scores above 93% (S1 Fig). Barcode rank plots clearly distinguished high-quality cells from background noise based on UMI count distribution(S2 Fig). A median of 389 to 752 genes per cell and sequencing saturation < 92% confirmed robust coverage across samples. Quality control metrics confirmed high data quality, with most cells expressing hundreds to thousands of genes and < 20% mitochondrial content (Fig 3A and 3B). After this, 22,672 cells were retained. Unsupervised clustering identified 19 clusters (Fig 3C and 3D). Guided by marker expression and reference-based annotation, these clusters were annotated and consolidated into 16 major cell populations (Fig. 3E), including T cells, NK cells, macrophages, Kupffer cells, hepatic stellate cells, hepatocytes, and other immune and parenchymal populations. Fig. 3F displays the top marker genes of clusters 0–8, while the remaining annotated populations are provided in S3 Fig.

**Fig 3.**
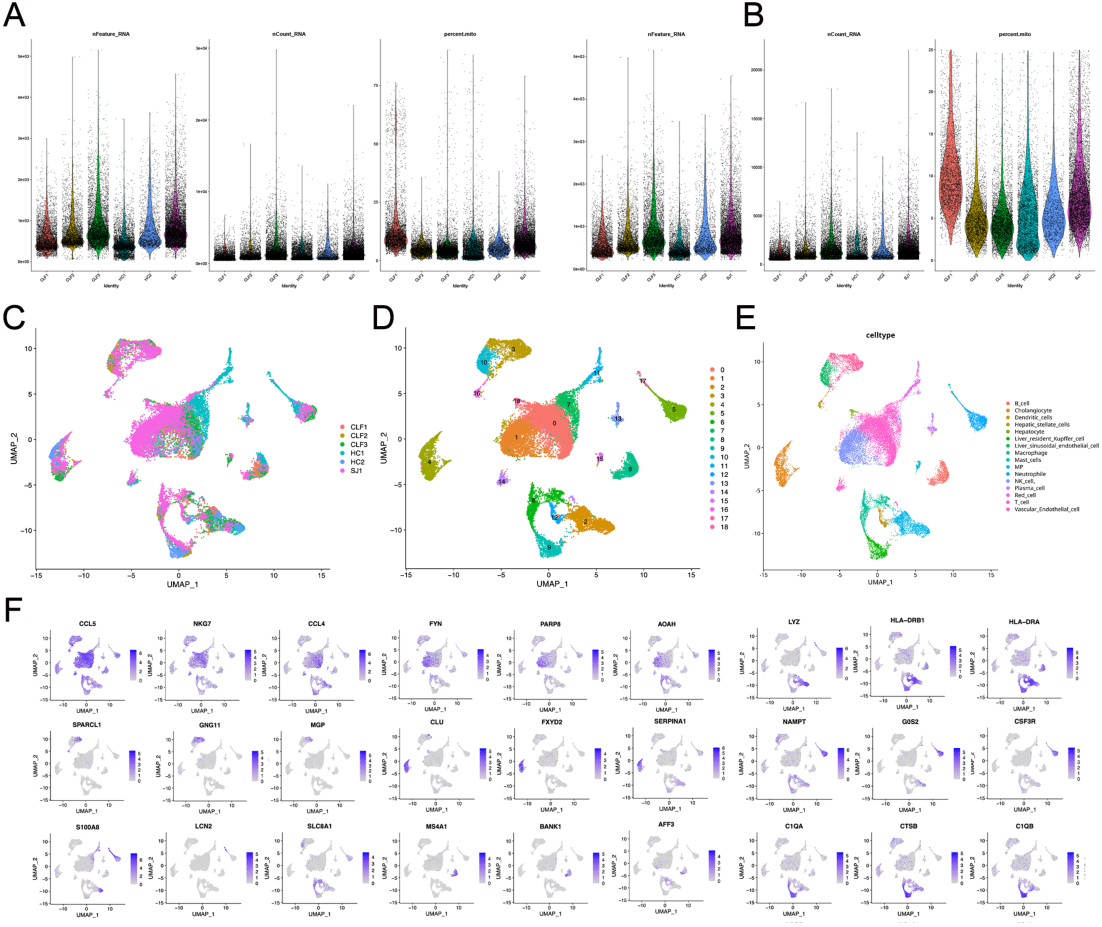
Single-cell transcriptomic profiling and cell type annotation of liver samples. (A–B) Violin plots of QC metrics before and after filtering, showing gene counts, UMI counts, and mitochondrial content. (C) UMAP of cells by sample origin. (D) UMAP of 19 clusters. (E) Refined UMAP of 16 major cell types. (F) Top three marker genes are shown for clusters 0–8.

### High-dimensional weighted gene co-expression network analysis (HD-WGCNA) and enrichment analysis

Using scale-free topology, β = 3 was chosen for network construction (Fig 4A). HD-WGCNA identified eight modules from 3,432 genes (Fig 4B). The turquoise module, containing 3,321 genes, showed the strongest correlation with SJ samples (r = 0.31, p < 0.05; Fig 4C) and most related with T cells (r = 0.12, p < 0.05; Fig 4D). GO and KEGG analyses highlighted protein phosphorylation, cytosolic localization, kinase activity, and the T cell receptor signaling pathway (Fig 4E and 4F).

**Fig 4.**
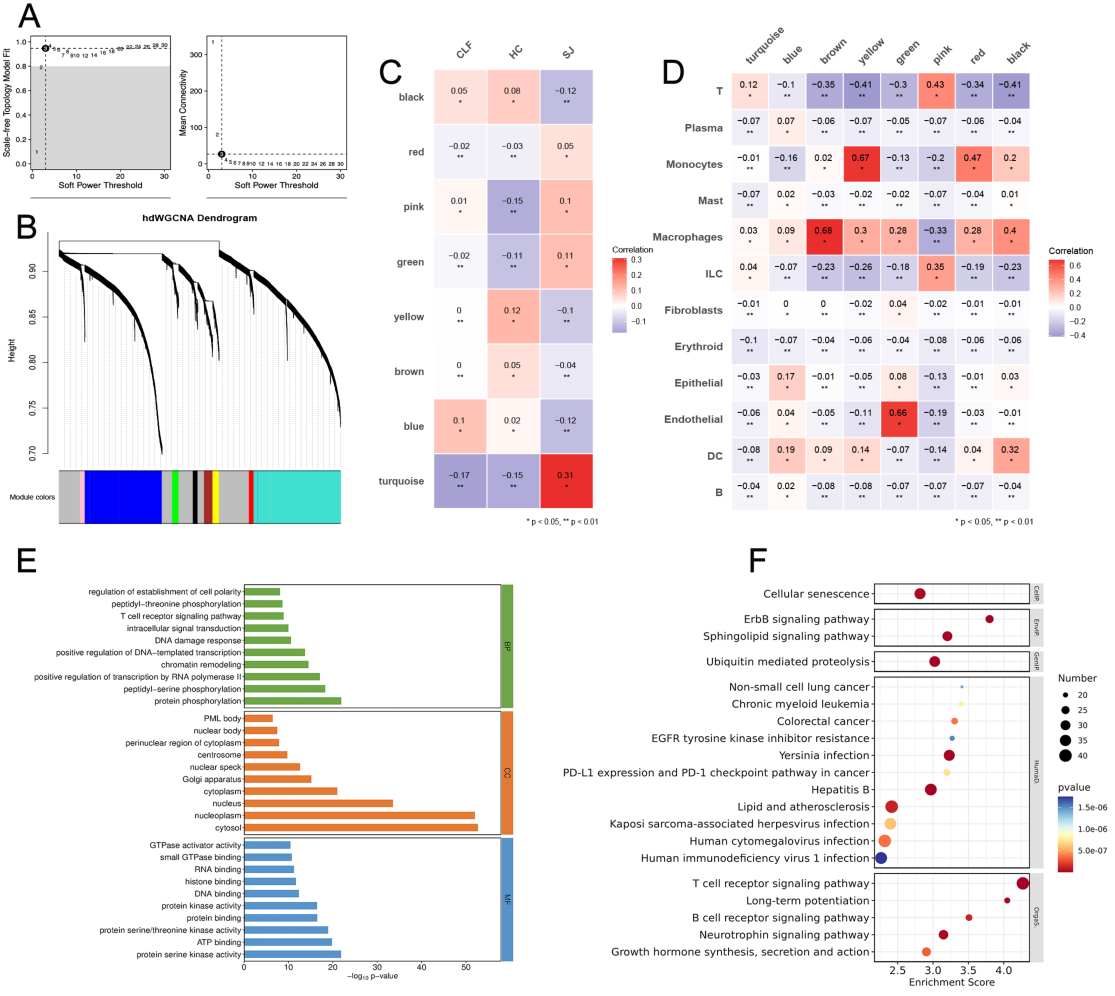
HD-WGCNAmodule identification and functional characterization.(A) Soft-thresholding power selection.(B) Gene dendrogram and module assignment. (C) Module–trait correlation heatmap.(D) Module–cell type correlation heatmap. (E) GO enrichment barplot.(F) KEGG enrichment dotplot.

### DEGs

From the SJ vs. HC comparison, 1,769 SJ-specific DEGs were obtained after excluding CLF-overlapping genes (Fig 5A). Intersecting these with SJ vs. CLF DEGs (1,187) and the turquoise module (1,493) yielded 219 SSLF-specific genes (Fig 5B). GO analysis highlighted small GTPase-mediated signaling, cytosol localization, and guanyl-nucleotide exchange activity (Fig 5C). KEGG analysis identified the T cell receptor signaling pathway as the top-enriched term (Fig 5D).

**Fig 5.**
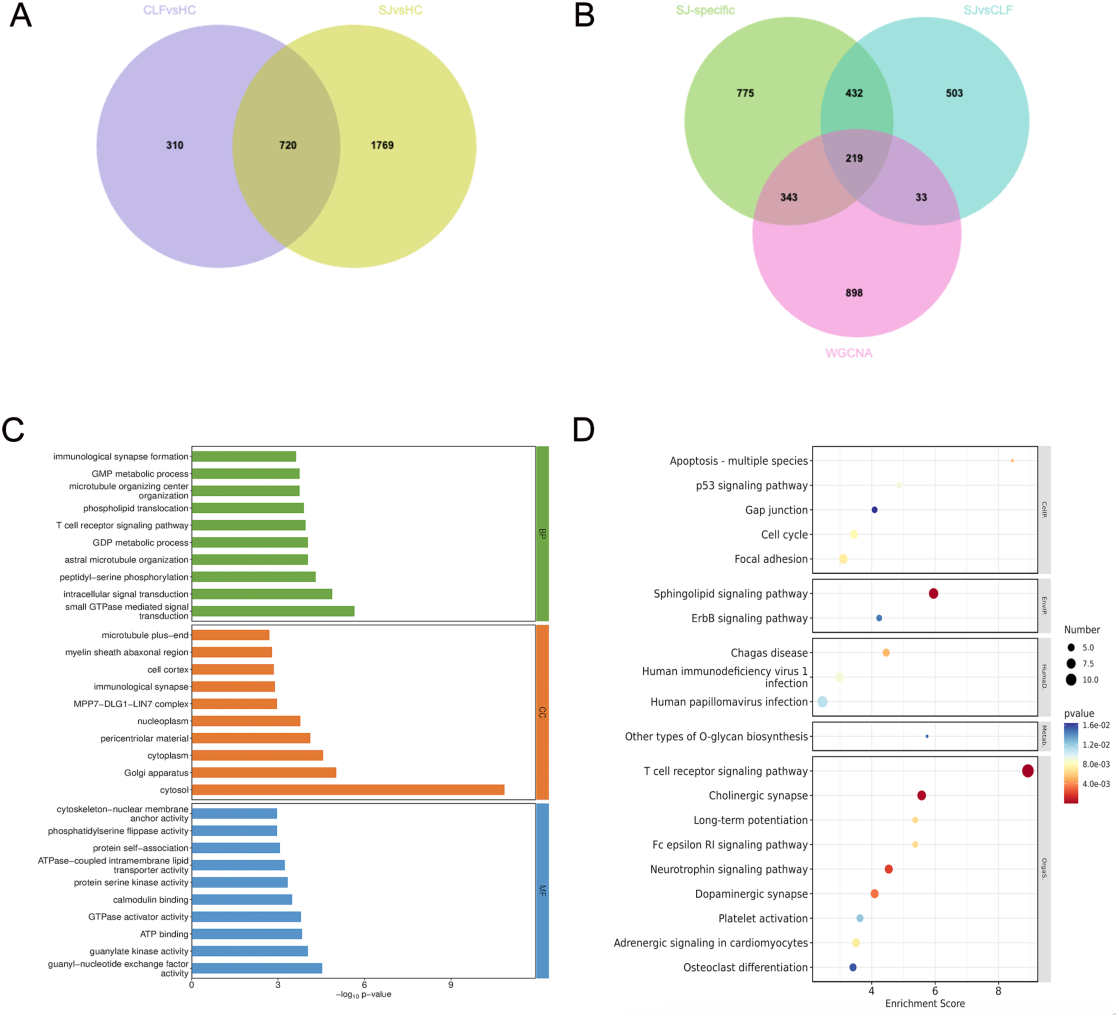
Differentially expressed genes analysis. (A) Venn gram of DEGs CLF vs HC and DEGs SJ vs HC.(B) Venn gram of SJ-specific DEGs, DEGs SJ vs CLF and Terquoise module of HD-WGCNA. (C) GO enrichment barplot. (D) KEGG enrichment dotplot.

### PPI, MCODE and hub gene identification

A total of 152 SSLF candidate genes were mapped into a PPI network (STRING, score ≥ 0.4) (Fig 6A). MCODE clustering revealed three significant modules (Fig 6B) and centrality analysis identified four hub candidates, including AKT3, FYN, BCL2 and ATM (Fig 6C). The overlap between these four hub candidates and modules yielded AKT3, FYN, and BCL2 as SSLF-related hub genes (Fig 6F). GO terms of the full PPI network were enriched in cytosol and nucleoplasm, while KEGG pathways highlighted T cell receptor and sphingolipid signaling (Fig 6D and 6E). Modules-specific enrichment further emphasized immune-related functions, including T cell receptor signaling and immunological synapse pathways (Fig 6G and 6H). These findings consistently point to T cell-related signaling as a central theme in the network.

**Fig 6.**
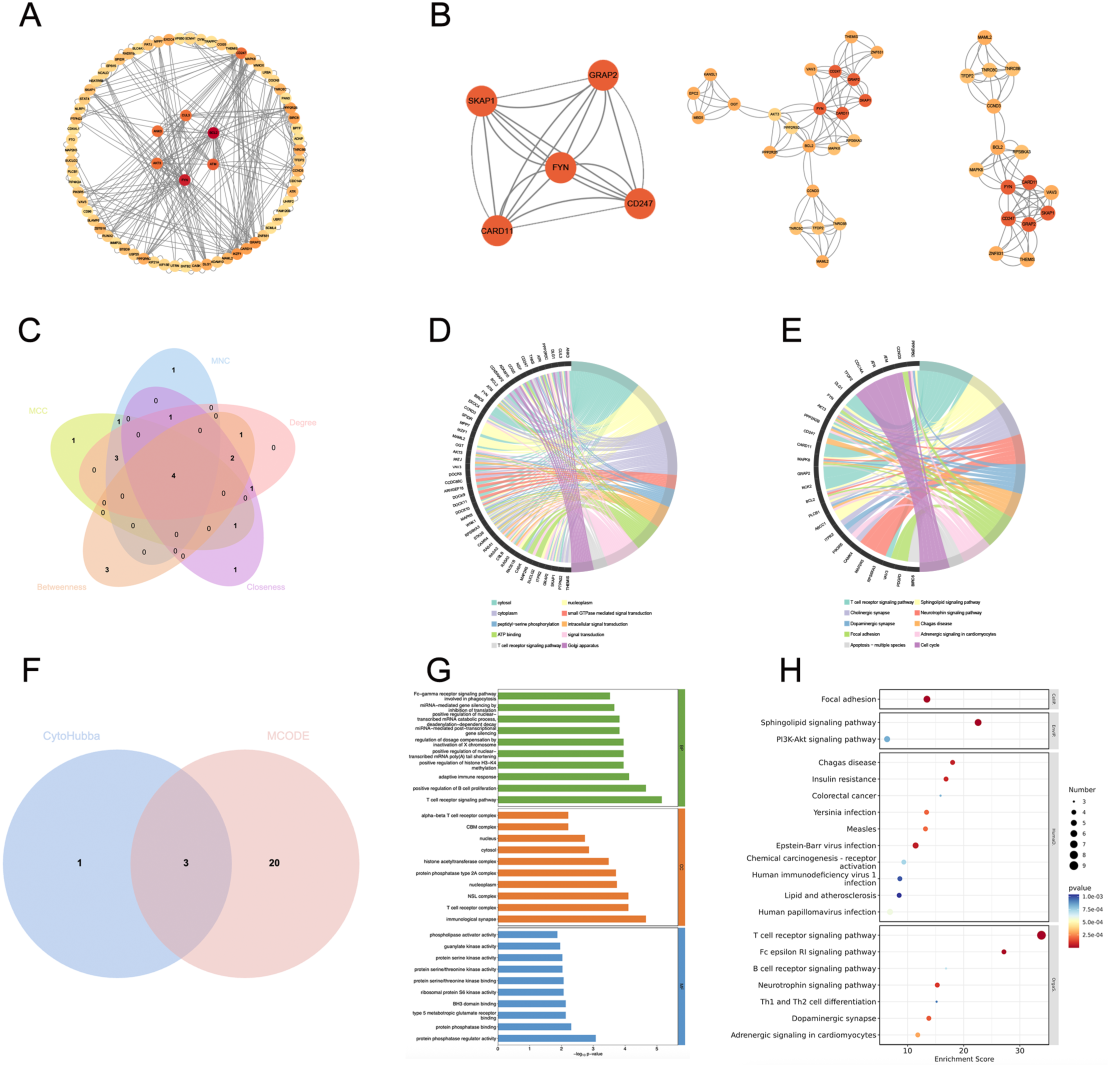
PPI, MCODE analysis and hub gene identification. (A) PPI network of SSLF candidate genes. (B) Top three MCODE modules. (C) Venn diagram of hub genes identified by five CytoHubba algorithms. (D–E) GO and KEGG enrichment of full PPI network. (F) Overlap of CytoHubba hub genes with MCODE modules. (G-H) GO and KEGG enrichment of module genes.

### Antigen screening

From UniProt, 29,870 *S. japonicum* antigens were retrieved, including 52 reviewed entries(S1 Table). GO annotation identified 25 antigens mapped to at least two categories(S2 Table); among them, 14 were annotated in all three(BP, CC, MF) and 11 in two categories(S4 Fig). Most were localized to nucleus or cytoplasm, with only PPIB_SCHJA containing a transmembrane helix and signal peptide, and CYSP_SCHJA carrying a signal peptide. BLAST analysis against the human proteome identified six low-homology candidates (< 30% identity): DRE2_SCHJA, GST28_SCHJA, ANX13_SCHJA, SORCN_SCHJA, MYSP_SCHJA, and CIAO1_SCHJA (Table 1).

**Table 1.** BLAST-based screening and predicted features of Schistosoma japonicum antigens.

| Antigen | TMHMM - 2.0 | Signal IP | DeepLoc - 2.0 | BLAST (%) |
| --- | --- | --- | --- | --- |
| PPIE_SCHJA | 0 | Nucleus | Nucleus | 62.5% |
| PPIB_SCHJA | 1 | 0.970780 | Endoplasmic reticulum,<br>Signal peptide | 59.7% |
| HGLB_SCHJA | 0 | 0.2436 | Extracellular | 39.6% |
| DRE2_SCHJA | 0 | 0 | Nucleus | 29.1% |
| PURA_SCHJA | 0 | 0 | Cytoplasm | 53.8% |
| ENO_SCHJA | 0 | 0 | Cytoplasm | 71.2% |
| KYNU_SCHJA | 0 | 0 | Cytoplasm | 38.6% |
| TPIS_SCHJA | 0 | 0 | Cytoplasm, Mitochondrion | 63.1% |
| RL24_SCHJA | 0 | 0 | Cytoplasm, Mitochondrion | 56.8% |
| RS3A_SCHJA | 0 | 0 | Cytoplasm | 59% |
| RSSA_SCHJA | 0 | 0 | Cytoplasm | 46.8% |
| CH10_SCHJA | 0 | 0 | Mitochondrion | 64% |
| ATAT_SCHJA | 0 | 0 | Nucleus | 32.1% |
| EIF3M_SCHJA | 0 | 0 | Cytoplasm, Nucleus | 32.5% |
| GST28_SCHJA | 0 | 0 | Cytoplasm | 28.8% |
| GST26_SCHJA | 0 | 0 | Cytoplasm | 38.5% |
| CYSP_SCHJA | 0 | 0.9986 | Extracellular<br>Signal peptide | 37.7% |
| CDO_SCHJA | 0 | 0 | Cytoplasm | 37.6% |
| HSP70_SCHJA | 0 | 0 | Cytoplasm | 66.5% |
| ANX13_SCHJA | 0 | 0 | Cytoplasm, Nucleus | 22.3% |
| FABP_SCHJA | 0 | 0 | Cytoplasm | 45% |
| TCTP_SCHJA | 0 | 0 | Cytoplasm, Nucleus | 45.9% |
| SORCN_SCHJA | 0 | 0 | Cytoplasm | 27.1% |
| MYSP_SCHJA | 0 | 0 | Cytoplasm | 18.4% |
| CIAO1_SCHJA | 0 | 0 | Cytoplasm, Nucleus | 20.9% |

### B-cell epitope identification

Among the six candidates, only DRE2_SCHJA carried a conserved B-cell epitope (EATEVED, 222–228) predicted as antigenic (Vaxijen = 0.8567), non-allergenic and non-toxic (Table 2). Although antigenic epitopes were present in GST28_SCHJA, ANX13_SCHJA, SORCN_SCHJA, and MYSP_SCHJA, they carried potential allergenicity. No antigenic B-cell epitopes were detected in CIAO1_SCHJA.

**Table 2.**
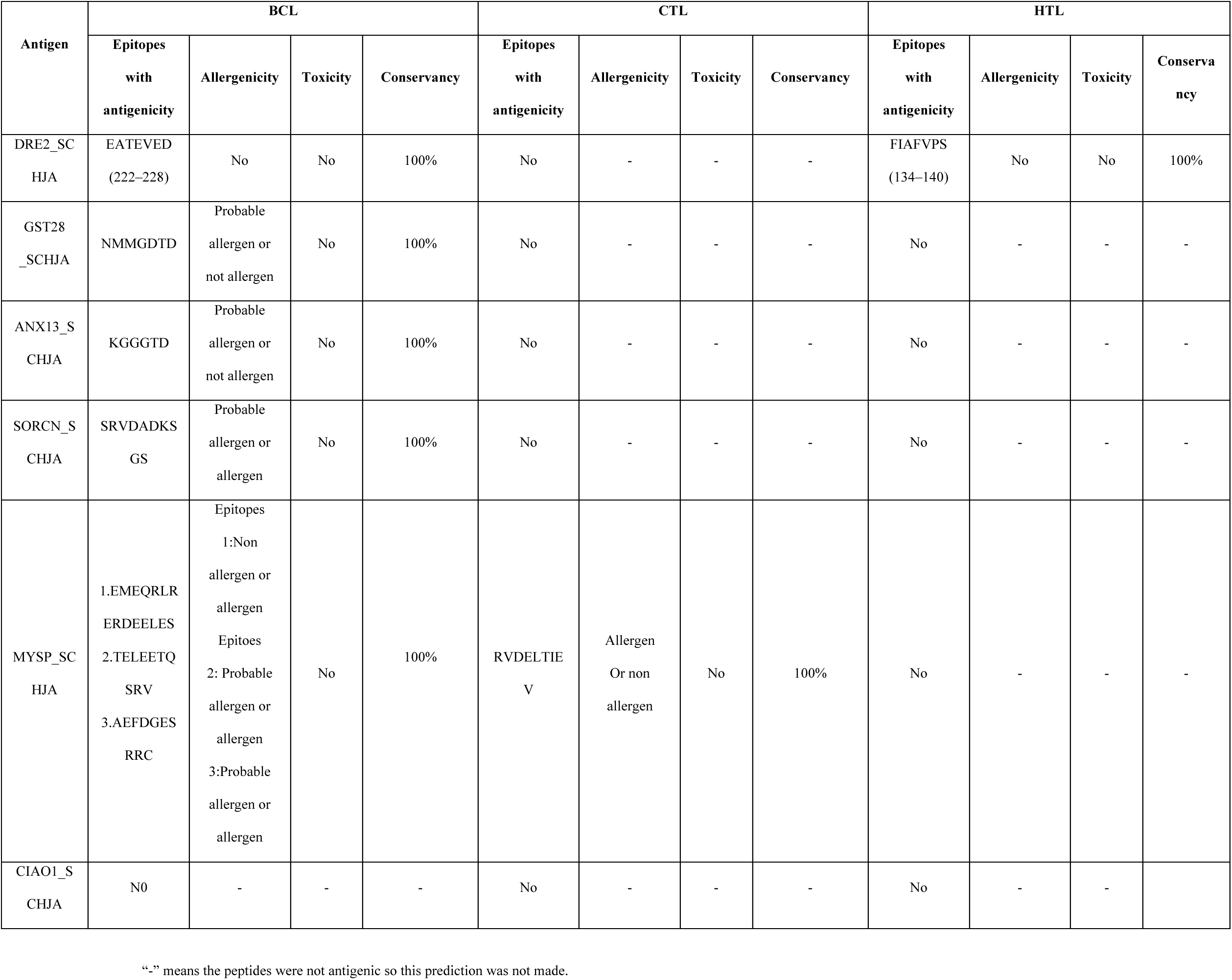
Predicted antigenicity, allergenicity, and toxicity of BCL, CTL, and HTL epitopes.

| Antigen | BCL |  |  |  | CTL |  |  |  | HTL |  |  |  |
| --- | --- | --- | --- | --- | --- | --- | --- | --- | --- | --- | --- | --- |
|  | Epitopes<br>with<br>antigenicity | Allergenicity | Toxicity | Conservancy | Epitopes<br>with<br>antigenicity | Allergenicity | Toxicity | Conservancy | Epitopes<br>with<br>antigenicity | Allergenicity | Toxicity | Conservancy |
| DRE2_SC<br>HJA | EATEVED<br>(222–228) | No | No | 100% | No | - | - | - | FIAFVPS<br>(134–140) | No | No | 100% |
| GST28<br>_SCHJA | NMMGDTD | Probable<br>allergen or<br>not allergen | No | 100% | No | - | - | - | No | - | - | - |
| ANX13_S<br>CHJA | KGGGTD | Probable<br>allergen or<br>not allergen | No | 100% | No | - | - | - | No | - | - | - |
| SORCN_S<br>CHJA | SRVDADKS<br>GS | Probable<br>allergen or<br>allergen | No | 100% | No | - | - | - | No | - | - | - |
| MYSP_SC<br>HJA | 1.EMEQRLR<br>ERDEELES<br>2.TELEETQ<br>SRV<br>3.AEFDGES<br>RRC | Epitopes<br>1:Non<br>allergen or<br>allergen<br>Epitopes<br>2: Probable<br>allergen or<br>allergen<br>3:Probable<br>allergen or<br>allergen | No | 100% | RVDELTIE<br>V | Allergen<br>Or non<br>allergen | No | 100% | No | - | - | - |
| CIAO1_S<br>CHJA | N0 | - | - | - | No | - | - | - | No | - | - | - |

“-” means the peptides were not antigenic so this prediction was not made.

### CTL epitope identification

Only MYSP_SCHJA contained a strong candidate (RVDELTIEV) with high antigenicity and no predicted toxicity, but inconsistent allergenicity across tools restricted its prioritization. None of the other antigens carried acceptable CTL epitopes (Table 2).

### HTL epitope identification

DRE2_SCHJA was the only antigen predicted to contain an HTL epitope (FIAFVPS, residues 134–140) fulfilling all criteria of antigenicity, non-allergenicity, non-toxicity, and full conservancy, whereas no HTL epitopes were identified in the other candidates (Table 2).

### Antigen selection

DRE2_SCHJA met all secondary selection criteria, exhibiting both B-cell and HTL epitopes that were consistently predicted to be antigenic, non-allergenic, non-toxic, and fully conserved. It was therefore selected as the only candidate for further study.

### Immune simulation of antigen

As shown in Fig. 7, DRE2_SCHJA induced a rapid and coordinated immune response. B-cell kinetics showed a sharp increase in antigen-internalizing and -presenting subsets within 5 days, followed by sustained activation for over 30 days, indicative of long-lived humoral potential (Fig 7A). Serological analysis revealed peak total IgM + IgG titers at day 15, with IgM dominating the early phase and IgG1/IgG2 levels rising steadily thereafter, consistent with class switching and durable antibody production (Fig 7B). Helper T-cell responses peaked at day 10 and remained near maximal, while memory Th subsets were established early and persisted throughout the 30-day period (Fig 7C). These findings suggest that the vaccine has the potential to induce long-lasting immune protection. Cytokine profiling indicated a Th1-skewed response, with IFN-γ and IL-12 peaking at day 7, accompanied by elevated TGF-β and IL-10, suggesting concurrent immune regulation (Fig 7D). Innate immune activation was evidenced by early dendritic cell and macrophage activation and antigen presentation (S5 and S6 Fig), providing the foundation for effective adaptive immunity. Together, DRE2_SCHJA might elicit a sustained immune response with both effector and regulatory features, supporting its potential as a vaccine candidate.

**Fig 7.**
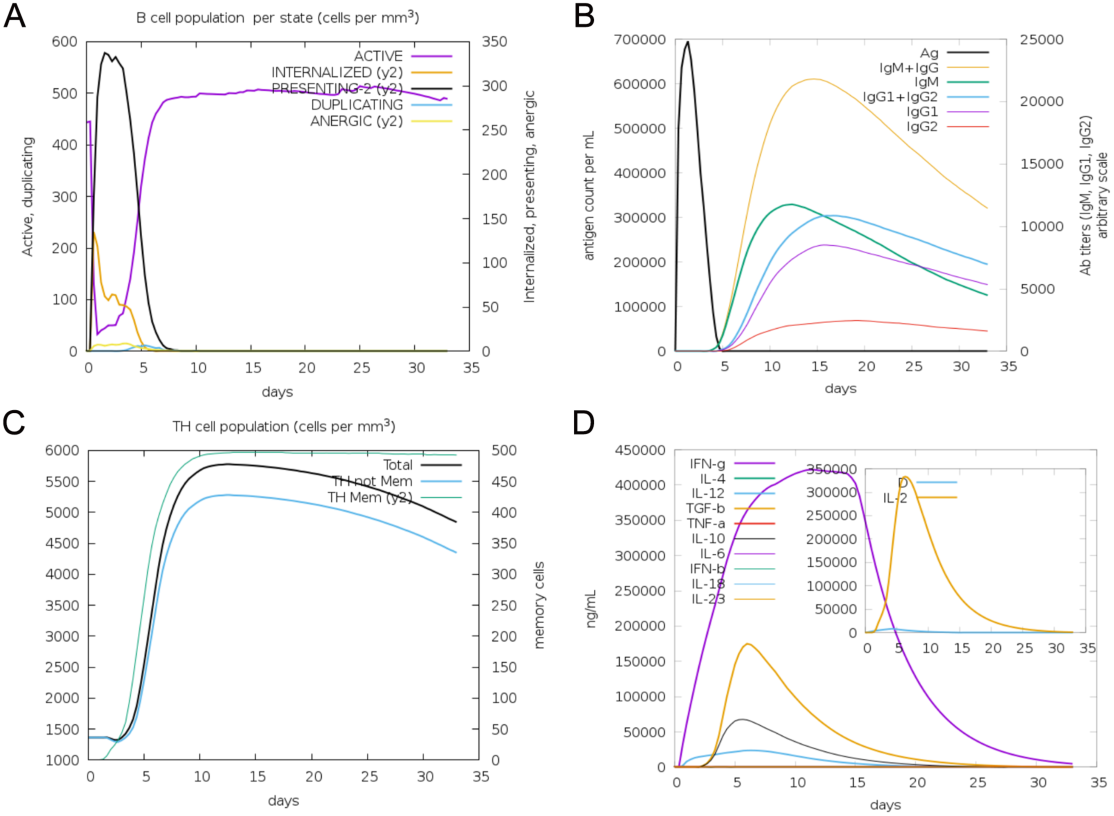
Immune simulation induced by DRE2_SCHJA in the C-ImmSim servers. (A) Expansion of active B cells (purple) following antigen stimulation. (B) Antigen clearance and total IgM + IgG production (orange) over time. (C) Induction of memory T helper cells (Green) following vaccination. (D) Cytokine release dominated by IFN-γ (purple) and IL-2 (yellow) peaks after immunization.

### First protein-protein docking

Protein-protein docking was performed using ClusPro to assess the interactions between DRE2_SCHJA and three host hub proteins: BCL2 (PDB ID: 8HTS), FYN (PDB ID: 4U1P), and AKT3 (PDB ID: 2X18). The DRE2_SCHJA–BCL2 complex showed a docking score of −264.258 and ΔG of −12.2 kcal/mol, supported by five hydrogen bonds and one salt bridge (Fig 8A). The DRE2_SCHJA–FYN complex achieved the most favorable interaction, with a docking score of −252.399 and a PRODIGY-predicted binding free energy of −14.9 kcal/mol, stabilized by six hydrogen bonds and one π–π interaction (Fig 8B). The DRE2_SCHJA–AKT3 complex had a docking score of −199.396 and ΔG of −13.8 kcal/mol, with three hydrogen bonds and four salt bridges (Fig 8C). Considering binding free energy, docking score, and interfacial interactions, DRE2_SCHJA–FYN formed the most stable complex, highlighting FYN as a potential central signaling node in SSLF.

**Fig 8.**
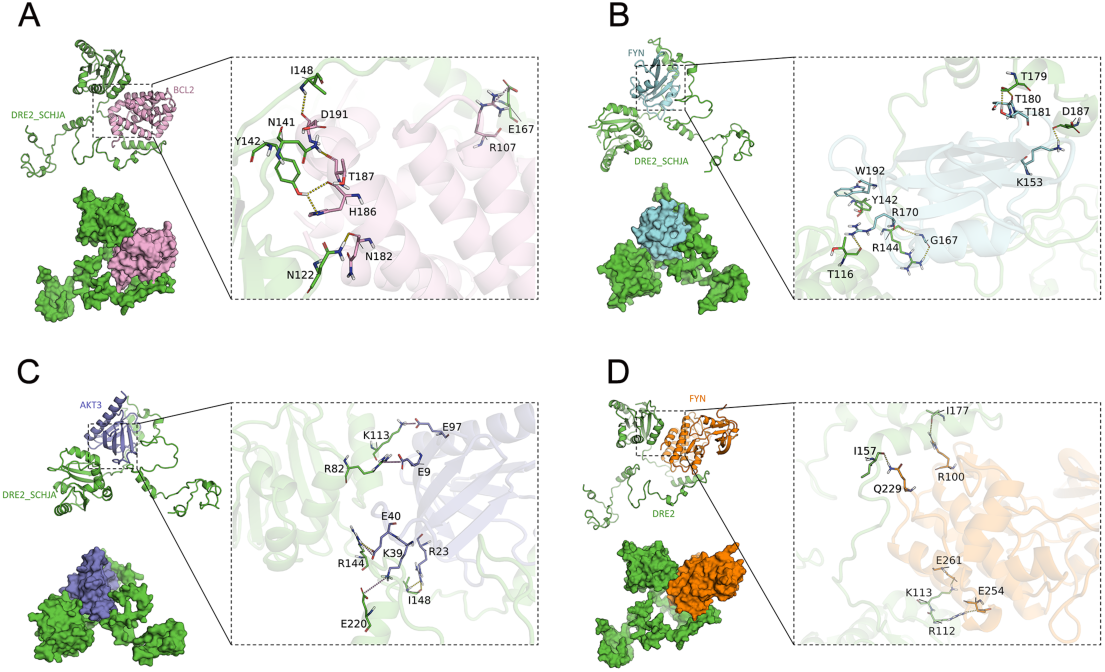
Protein–protein docking of antigen and host protein.(A) Docking of DRE2_SCHJA–BCL2. Green protein is DRE2_SCHJA, purple protein is BCL2. (B) Docking of DRE2_SCHJA–FYN (PDB ID: 4U1P). Green protein is DRE2_SCHJA, blue protein is FYN. (C) Docking of DRE2_SCHJA–AKT3. Green protein is DRE2_SCHJA,dark blue protein is AKT3. Yellow dashes represent hydrogen bonds, pink dashes indicate salt bridges, and blue dashes indicate π–π stacking interactions. (D) Docking of DRE2_SCHJA–FYN(PDB ID: 2DQ7). Yellow dashes represent hydrogen bonds.

### Molecular docking

Molecular docking using Maestro reproduced the previously reported binding energy and binding mode of Saracatinib to FYN (PDB ID: 2DQ7) reported by Dhanushya Gopal et al.[34]. The predicted docking score (−9.251 kcal/mol) closely matched the reported value. In our model, the ligand formed hydrogen bonds with Lys87 and Asp96, supported by hydrophobic contacts with Ile80, Tyr84, and Met85, and electrostatic stabilization by Glu83 (Fig 9A). Consistent with prior findings, Saracatinib occupied the ATP-binding pocket of FYN in the 3D model, stabilized through hydrogen-bonding and hydrophobic interactions (Fig 9B). The surface representation further illustrated its spatial orientation within the binding cavity (Fig 9C), reinforcing the reliability of the docking predictions.

**Fig 9.**
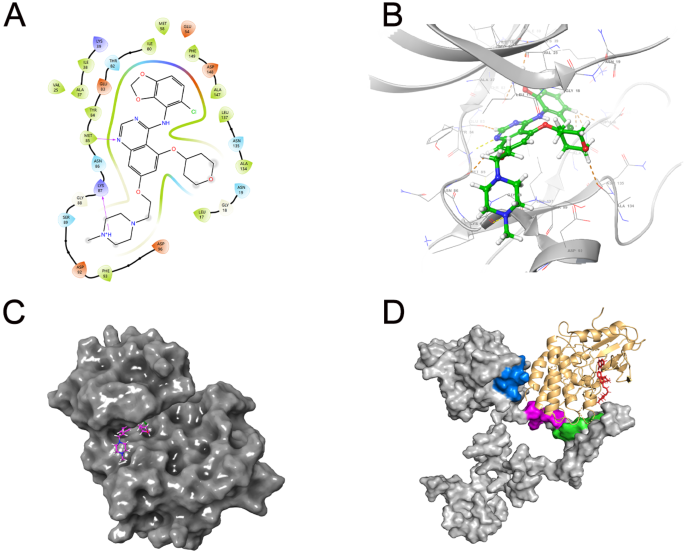
Docking validation of FYN with Saracatinib and DRE2 binding modes (A) 2D interaction map showing hydrogen bonds, hydrophobic and electrostatic contacts. (B) 3D binding conformation of Saracatinib (green sticks) in the ATP-binding pocket. (C) Surface view of FYN with Saracatinib (magenta sticks) located in the binding cavity. (D) Structural comparison of DRE2–FYN and Saracatinib–FYN complexes based on the FYN crystal structure (PDB ID: 2DQ7). FYN is shown in brown, DRE2_SCHJA in grey, and Saracatinib in red. DRE2_SCHJA binding regions are highlighted in blue, purple, and green.

### Secondary protein-protein docking

Docking of DRE2_SCHJA with FYN (PDB ID: 2DQ7) yielded a docking score of −122.483 kcal/mol and a predicted binding free energy of −9.3 kcal/mol. The interface was stabilized by four hydrogen bonds involving FYN residues Arg100, Arg112, Lys113, and Ile157 and DRE2 residues Ile177, Glu254, Glu261, and Gln229 (Fig 8D), indicating a stable complex.

### Visualization

Structural alignment demonstrated that DRE2_SCHJA interacts with residues located in the SH2 domain of FYN (PDB ID: 2DQ7), consistent with the docking interface identified in FYN (PDB ID: 4U1P) (Fig 8B and 8D). In contrast, Saracatinib binds within the ATP–binding pocket (Fig 9A). The superimposed structural model confirmed that the docking sites of DRE2 and Saracatinib occupy distinct, non-overlapping regions of FYN (Fig 9D).

### Motif prediction

Motif analysis predicted the pY-containing motifs in DRE2_SCHJA (S1 File). In the FYN (PDB ID: 4U1P)–DRE2_SCHJA complex, residues R144, D187, and Y142 fell within the SH2 motif, with T116 positioned nearby. This suggested a canonical SH2–pY recognition. In contrast, the FYN (PDB ID: 2DQ7) complex involved residues located outside the motif, indicating a non-canonical binding mode.

## Discussion

SSLF is the most serious chronic complication of *S. japonicum*, causing heavy disease burden, high medical costs, and poor quality of life. Current Schistosomiasis vaccine candidates in clinical development mainly aim to reduce parasite burden or egg output[37], but rarely address prevention or treatment of chronic pathology such as SSLF. In China, *S. japonicum* infection has been well controlled, yet SSLF persists as a major health burden. Developing a targeted vaccine or therapy is therefore of particular importance, as preventing fibrosis would directly reduce long-term morbidity and disability. RV uses genome-derived immunoinformatics to predict antigens with strong immunogenicity and good safety[38]. This approach speeds up candidate discovery, reduces reliance on animal experiments, and overcomes the slow and costly screening of traditional vaccine development[39]. However, Current RV pipelines are primarily designed to identify antigens that elicit strong immune responses, but they seldom incorporate host-pathology context when prioritizing candidates[13]. In this study, we addressed this gap by focusing on SSLF and developing an in silico pipeline integrating immunoinformatics, safety prediction, and host-pathology insights. This approach might enable the early identification of antigens with both protective potential and the capacity to influence SSLF progression. Using this strategy, we prioritised DRE2_SCHJA, which met immunogenicity and safety criteria and showed the strongest predicted interaction with FYN, a fibrosis-associated hub protein. This highlights the value of incorporating pathology-oriented criteria into RV workflows to guide targeted vaccine design for SSLF.

Among the fibrosis-related hub genes identified, FYN emerged as a key upstream regulator. As a member of the Src family tyrosine kinases, it acts early in T-cell receptor (TCR) signaling, linking antigen recognition to hepatic stellate cell (HSC) activation and extracellular matrix (ECM) remodeling[40]. Beyond fibrogenesis, FYN also participates in immune-checkpoint-regulated pathways and contributes to T-cell dysfunction under chronic stimulation[41]. Overall, these findings indicate that FYN may act as a node integrating pro-fibrotic and immune regulatory processes. In contrast, BCL2 and AKT3 were positioned as downstream effectors. BCL2 regulates apoptosis, while AKT3 is involved in metabolic control, together supporting cell survival and functional adaptation in the fibrotic liver[42, 43]. Their position downstream of FYN shows that there’s a hierarchical signaling network where FYN is at the center of regulation. Evidence from pharmacological studies supports this framework. The Src-family inhibitor Saracatinib (AZD0530) has been shown to attenuate liver fibrosis in vivo by blocking the FYN–FAK–N–WASP axis [44]. This reinforces the role of FYN as a critical control node in fibrosis-related signaling. From a translational perspective, FYN is an actionable target.

Among the parasite antigens analyzed, DRE2_SCHJA was prioritized as the lead candidate. It encodes an Anamorsin homolog that functions in the cytosolic iron–sulfur (Fe–S) cluster assembly pathway, which is essential for Fe–S protein maturation[45]. In *S. japonicum*, this pathway supports mitochondrial metabolism, redox homeostasis, and reproductive capacity[46]. As an upstream regulator of Fe–S cluster assembly, DRE2_SCHJA may serve as a strategic intervention point in parasite metabolism. Fe–S-dependent proteins such as SjSDISP are indispensable for egg production, and their disruption reduces egg burden and hepatic fibrosis in murine models[47, 48], highlighting the close link between Fe–S metabolism and pathology. Other Fe–S-associated proteins, such as SjCIAP, further support the essentiality of this pathway. Beyond Schistosomiasis, Fe–S clusters are increasingly recognized as transcriptionally regulated mediators of diverse pathological processes, underscoring their broader therapeutic relevance[49]. These findings show their emerging therapeutic relevance and support the idea that targeting Fe–S metabolism is a viable and strategically important approach. Egg deposition remains the key driver of hepatic fibrosis[46]. Thus, targeting Fe–S cluster assembly could impair parasite reproduction, limit granuloma formation, and slow fibrotic progression. However, potential functional redundancy within parasite metabolic networks may compensate for DRE2 inhibition. Further experimental validation is needed to confirm whether disruption of Fe–S metabolism can achieve meaningful antifibrotic benefits while ensuring antigen accessibility in vivo.

Alongside its metabolic role, DRE2_SCHJA was also predicted to shape immune responses in ways that could help limit liver fibrosis. Predicted IL-2 induction could promote effector T-cell expansion and macrophage activation[50], therefore enhancing the clearance of pro-fibrotic stimuli and accelerating the removal of inflammatory triggers. Increased IFN-γ can inhibit hepatic stellate cell activation and suppress collagen synthesis via STAT1 signaling[51], thereby directly attenuating fibrogenesis. Besides, sustained B-cell activity may contribute to IL-10 production, which downregulates inflammatory chemokines and reduces monocyte recruitment[52]. Class-switched IgG could further restrict egg-antigen exposure and limit chronic immune stimulation[53]. The cytokine environment also indicates a balance between pro- and antifibrotic forces: TGF-β may promote early fibrosis but later cooperate with IL-10 to restrain T cell–driven inflammation[54]. Together with elevated IFN-γ and IL-12, this pattern reflects a Th1-skewed response that counteracts the profibrotic Th2 bias characteristic of Schistosomiasis[55]. Collectively, these predicted features suggest that DRE2_SCHJA might mobilize both effector and regulatory mechanisms to favor an antifibrotic state, supporting its candidacy as a promising vaccine target for SSLF.

Docking analysis identified FYN as the strongest interactor of DRE2_SCHJA, consistent with its established role in immune activation and fibrotic signaling. Two distinct SH2 binding modes were predicted. In the pY-proximal mode (PDB ID: 4U1P), DRE2_SCHJA contacted residues Y142 and R144 located within the canonical phosphotyrosine recognition pocket. Such positioning suggests potential competition with phosphorylated adaptor proteins[56], which might disrupt SH2–pY interactions[57, 58]. This disruption could in turn weaken early T-cell receptor signaling and may attenuate downstream pathways such as PI3K–AKT, JAK–STAT, and Src–YAP[59–61]. As a result, T-cell hyperactivation might be reduced, cytokine secretion may be limited, and hepatic stellate cell activation could be suppressed, thereby slowing the progression of fibrosis[62, 63]. In contrast, the pY-distal mode (PDB ID: 2DQ7) involved residues such as Arg100 and Lys113, located outside the canonical pocket. This non-canonical binding pattern is consistent with evidence that SH2 domains can act as allosteric regulators[64, 65]. Engagement at these distal sites may perturb the SH2–SH3 linker or the SH2-kinase interface[66]. Both regions are structural hubs that influence FYN conformational activation. Distal binding could therefore weaken conformational activation by altering cross-domain communication, and may modulate downstream pathways without directly competing for phosphotyrosine motifs[67, 68]. These SH2-based interactions differ from ATP-competitive inhibition, as observed with Saracatinib, which targets the catalytic site of FYN[69]. Instead, DRE2_SCHJA appears to act through adaptor interference or interdomain modulation. In summary, these two predicted modes suggest alternative ways in which a parasite-derived antigen may influence FYN signaling, providing potential routes for modulating fibrosis-related pathways in SSLF.

In conclusion, our docking analyses suggest that the parasite-derived antigen DRE2_SCHJA fulfills key criteria for vaccine development and may modulate host fibrotic signaling through SH2–mediated interactions with FYN. While these results are based on computational predictions and require experimental validation, they show the feasibility of pathology-oriented vaccination strategies. This approach is especially important for SSLF, where fibrosis control remains an unmet need. By integrating host–parasite single-cell transcriptomics with reverse vaccinology and structure-based modeling, this study provides a framework for identifying SSLF antigens and advancing targeted vaccine development for Schistosomiasis.

## Conclusion

DRE2_SCHJA emerges as a unique pathology-oriented vaccine candidate with potential dual effects: it may help prevent the onset of SSLF and modulate host fibrotic signaling. While based on computational predictions, this study demonstrates a methodological innovation by integrating reverse vaccinology with host-parasite single-cell data. This integrated approach enables the early identification of antigens in a rapid, efficient, and cost-effective manner, providing a framework to guide pathology-oriented vaccine development for Schistosomiasis and other neglected tropical diseases.

## Limitation

This study has several limitations. First, due to the scarcity of *S. japonicum* samples, the analysis relied on a public single-cell RNA-sequencing dataset from livers co-infected with *S. japonicum* and HBV. While co-infection is common, it could potentially introduce HBV-related confounding signals, although multi-step filtering was applied to minimize this effect. Second, the single-cell analysis focused on T cells, which play a key role in fibrogenesis, but other immune and stromal populations warrant further investigation in future studies. Third, antigen selection was restricted to curated proteins to ensure annotation reliability, which may have excluded novel candidates. Fourth, as DRE2_SCHJA is proposed here for the first time as a vaccine target, its functional relevance requires experimental validation, particularly during egg deposition and host invasion. Finally, while docking suggested stable binding between DRE2_SCHJA and FYN, its downstream impact on phosphorylation recognition and fibrotic signaling remains to be determined. Future work integrating in vitro validation, in vivo functional studies, and longitudinal vaccine trials will be essential to establish the translational potential of DRE2_SCHJA.

## Author contributions

Yang Qian Xu contributed to methodology, formal analysis, validation, visualization, and writing original draft. Dr. Wang Jue performed formal analysis related to molecular docking. Prof. Dr. Ming Ying Zi and Dr. Zhang Yu contributed to resources and data curation, providing the single-cell RNA sequencing data and depositing it in a public database. Prof. Dr. Wong Li Ping contributed to review & editing and provided continuous supervision during manuscript development. Dr. Lee Hai Yen was responsible for conceptualization, methodology, supervision, and funding acquisition, including providing overall technical support and methodological guidance. All authors reviewed and approved the final version of the manuscript.

## Supporting information

**S1 Fig. Data quality control**

**S2 Fig. Effective cell evaluation chart for all samples**

**S3 Fig. Top three marker genes of cluster 9-15**

**S4 Fig. Venn gram of the BP, CC and MF antigens**

**S5 Fig. Induction of dendritic cell population following vaccination**

**S6 Fig. Induction of macrophage population following vaccination**

**S1 Table. Fifty-two *S. japonicum* antigen entries**

**S2 Table. The overlapping antigens among BP, CC and MF**

**S1 File. Motif prediction of DRE2_SCHJA**

## Acknowledgement

We acknowledge Biozeron Co., Ltd. for performing the technical scRNA-seq data processing, and for ensuring methodological accuracy through result review and repeated analyses when necessary.

## Funding

This study was supported by the Research University Grant (RU006-2025G) and the MOSTI Strategic Research Fund (MOSTI003-2021SRF). The research was also partially supported through access to facilities funded by the Ministry of Higher Education, Malaysia, under the Higher Institution Centre of Excellence (HICoE) program (MO002-2019 & TIDREC-2023).

## Notes

### Competing Interest Statement

The authors have declared no competing interest.

